## Supplemental figures for "The propensity of fructose to induce metabolic dysfunction is dependent on the baseline diet, length of the dietary exposure, and sex of the mice"

**Supplementary tables**

**Table 1. Mouse primers used for real-time quantitative PCR.**

| Gene | Sequence (5' – 3') | Forward/Reverse |
|---|---|---|
| Acly | CAGCCAAGGCAATTTCAGAGC | Forward |
| | CTCGACGTTTGATTAACTGGTCT | Reverse |
| Acc1 | ACAGTGGAGCTAGAATTGGAC | Forward |
| | ACTTCCCGACCAAGGACTTTG | Reverse |
| Fasn | GGAGGTGGTGATAGCCGGTAT | Forward |
| | TGGGTAATCCATAGAGCCCAG | Reverse |
| Scd1 | CAGCCGAGCCTTGTAAGTTC | Forward |
| | GCTCTACACCTGCCTCTTCG | Reverse |
| Khk-c | AACTCCTGCACTGTCCTTTCCTT | Forward |
| | CCACCAGGAAGTCGGCAA | Reverse |
| Cpt1a | AGTGGCCTCACAGACTCCAG | Forward |
| | GCCCATGTTGTACAGCTTCC | Reverse |
| Khk-a | TTGCCGATTTTGTCCTGGAT | Forward |
| | CCTCGGTCTGAAGGACCACAT | Reverse |
| 18S | GTAACCCGTTGAACCCCATT | Forward |
| | CCATCCAATCGGTAGTAGCG | Reverse |
| Tbp | TGACTGCAGCAAATCGCTTGG | Forward |
| | ACCCTTCACCAATGACTCCTATG | Reverse |
| ACOX1 | ACTCGCAGCCAGCGTTATG | Forward |
| | AGGGTCAGCGATGCCAAAC | Reverse |
| ACOT1 | AGAGGAAGAGTTGGGCAGAG | Forward |

| | | |
|---|---|---|
| | TTCGTCCCAGCAGCAGCG | Reverse |

**Table 2. Antibodies used for Western Blot.**

| Name | Citation | Supplier | Cat no. |
|---|---|---|---|
| Rabbit polyclonal anti-ACLY | RRID: AB_2223744 | Cell Signaling Technology | 4332 |
| Rabbit monoclonal anti-ACC | RRID: AB_2219397 | Cell Signaling Technology | 3676 |
| Rabbit monoclonal anti-FASN | RRID: AB_2100796 | Cell Signaling Technology | 3180 |
| Rabbit polyclonal anti-SCD1 | RRID: AB_823634 | Cell Signaling Technology | 2438 |
| Rabbit monoclonal anti-KHK-A | | Signal way Antibody | 21708 |
| Rabbit monoclonal anti-KHK-C | | Signal way Antibody | 21709 |
| Mouse monoclonal anti-CPT1a | RRID: AB_11141632 | Abcam | ab128568 |
| Mouse monoclonal anti-Vinculin | RRID: AB_2272814 | Novus Biologicals | NB600-1293 |
| Rabbit polyclonal anti-OCTN2 | RRID: AB_2191406 | Proteintech | 16331-1-AP |
| Rabbit polyclonal anti-CACT | RRID: AB_10642001 | Proteintech | 19363-1-AP |

| | | | |
|---|---|---|---|
| Rabbit polyclonal anti-CPT2 | RRID: AB_2084849 | Abcam | ab71435 |
| Mouse monoclonal anti-ACADVL | RRID: AB_10609094 | Santa-Cruz | sc-271225 |
| Rabbit polyclonal anti-ACADL | RRID: AB_1859818 | Abcam | ab82853 |
| Mouse monoclonal anti-Actin | RRID: AB_476744 | Sigma | A5441 |
| Rabbit monoclonal anti-pAkt | RRID: AB_2315049 | Cell Signaling Technology | 4060 |
| Rabbit monoclonal anti-total Akt | RRID: AB_915783 | Cell Signaling Technology | 4691 |
| Rabbit monoclonal anti-pERK | RRID: AB_2315112 | Cell Signaling Technology | 4370 |
| Rabbit monoclonal anti-total ERK | RRID: AB_330744 | Cell Signaling Technology | 9102 |
| Mouse monoclonal anti-HADHA | RRID: AB_ 10862577 | Abcam | ab110302 |
| Mouse monoclonal anti-ACOX1 | RRID: AB_ 3075447 | Santa-Cruz | sc-517306 |
| Mouse monoclonal anti-ACOT1 | RRID: AB_10918465 | Santa-Cruz | sc-373917 |
